# An Artificial Intelligence Model for Translating Natural Language into Functional de Novo Proteins

**DOI:** 10.1101/2025.03.21.644400

**Authors:** Timothy P. Riley, Mohammad S. Parsa, Pourya Kalantari, Ismail Naderi, Kiana Azimian, Nemya Begloo, Oleg Matusovsky, Kooshiar Azimian, Kathy Y. Wei

## Abstract

Traditional protein design is fundamentally constrained by known sequences and folds. To break free from these limitations, we introduce a new alternative: designing proteins directly from plain-language specifications. To achieve this, we trained MP4, a transformer-based model that maps natural language prompts to protein sequences, on a dataset of 3.2 billion points and 138k tokens. In a benchmark of 96 prompts representing a wide array of functions and contexts, MP4 excelled by simultaneously improving on three key metrics: sequence realism, predicted fold quality, and alignment to the requested function. This high performance is particularly significant as it was achieved using only text as input which is a major departure from other models. Experimental validation confirmed our computational predictions: two de novo designs were experimentally shown to be both expressible and thermostable, with high-resolution crystallography (1.30 Å and 1.77 Å) ultimately revealing one to possess a paradigm-shifting novel fold. Functionally, the designs were also active, demonstrating both ATP binding and hydrolysis *in vitro*. This work demonstrates the realization of natural-language intent as functional proteins that express, crystallize, and catalyze. Although the underlying approach is still in early development with incomplete coverage and controllability, MP4 delivers a profound impact: it lowers the barrier to protein design and vastly expands the space for creative exploration in molecular programming.

## Introduction

Designing proteins from natural language prompts marks a paradigm shift from conventional template-based methods. Traditional approaches require a predefined molecular scaffold—a specific sequence or motif structure—which is then iteratively optimized for a target function(*1–8*). This template-first paradigm constrains the design process to local regions of the vast protein space, requiring the scientist to select a plausible starting point—for example, to identify a motif for “ATP hydrolysis” or a sequence likely to bind a given ligand—before optimization can begin. In contrast, text-based design aims to specify function and context up front, allowing the model to internalize possible starting sequences or structures during generation and thereby widen the accessible search space. In contrast, text-based generation aims to specify desired functional and structural properties a priori, tasking the model with learning a mapping from this high-level description to a plausible amino acid sequence and thereby unlocking a potentially much broader and more diverse design space.

Recent advances in machine learning have established text as a practical interface for molecular design. Inspired by large language models, text-to-protein generative models treat functional specifications —such as ‘ATP-binding enzyme’ or ‘secreted dimeric cytokine’— as input tokens and generate protein sequences as output. Early explorations by industry and academia, such as ESM3, ProGEN, and ProteinDT, have demonstrated the feasibility of this approach, while also exposing key challenges in controllability, prompt engineering, and the translation of linguistic intent into the biochemical and structural “grammar” of proteins(*9–15*). Instead of training new models directly, other approaches leverage large language models to interpret principles of biology and chemistry, but with the very similar goal of making it easier to translate words into functional molecules(*16*, *17*). The central premise is that text-based pipelines can be less constrained, more generalizable, and creative—broadening participation in protein design by lowering barriers for newcomers, while simultaneously empowering experts with new tools to explore the protein universe in ways that were previously inaccessible.

Here, we introduce the Molecular Programming model, version 4 (MP4): a transformer-based generative model that designs functional protein sequences from textual descriptions alone, requiring no initial sequence or structural input. We trained MP4 on a comprehensive dataset comprising 3.2 billion unique data points. This process utilized a unified vocabulary of 138k tokens to bridge natural-language descriptors with protein sequences. We evaluated MP4 by generating novel protein sequences from diverse textual prompts, analyzing their biophysical properties, and benchmarking against state-of-the-art models, such as ESM3 and Pinal(*9*, *10*). To confirm the real-world efficacy of our approach, a selection of computationally designed proteins were synthesized and experimentally validated. The results provide a clear proof-of-principle: MP4 generates novel proteins that successfully express, fold, crystallize, and exhibit catalytic activity. While this study marks a significant advance, we acknowledge that limitations in functional coverage, controllability, and interpretability remain active areas for future research.

## Results

### MP4 is a molecule programming model that transforms text into proteins

MP4 is a transformer-based, text-to-protein model that translates natural-language prompts into protein sequences aligned to input functions and properties (Fig. 1A). Unlike established template-driven workflows that begin from a known sequence or backbone, MP4 can take text as input and generate sequences directly, enabling intent-driven design from minimal specifications. The concept of designing straight from text is influenced and enabled by recent machine learning advances in natural language models and allows for more flexible inputs and more diverse outputs.

**Fig. 1.**
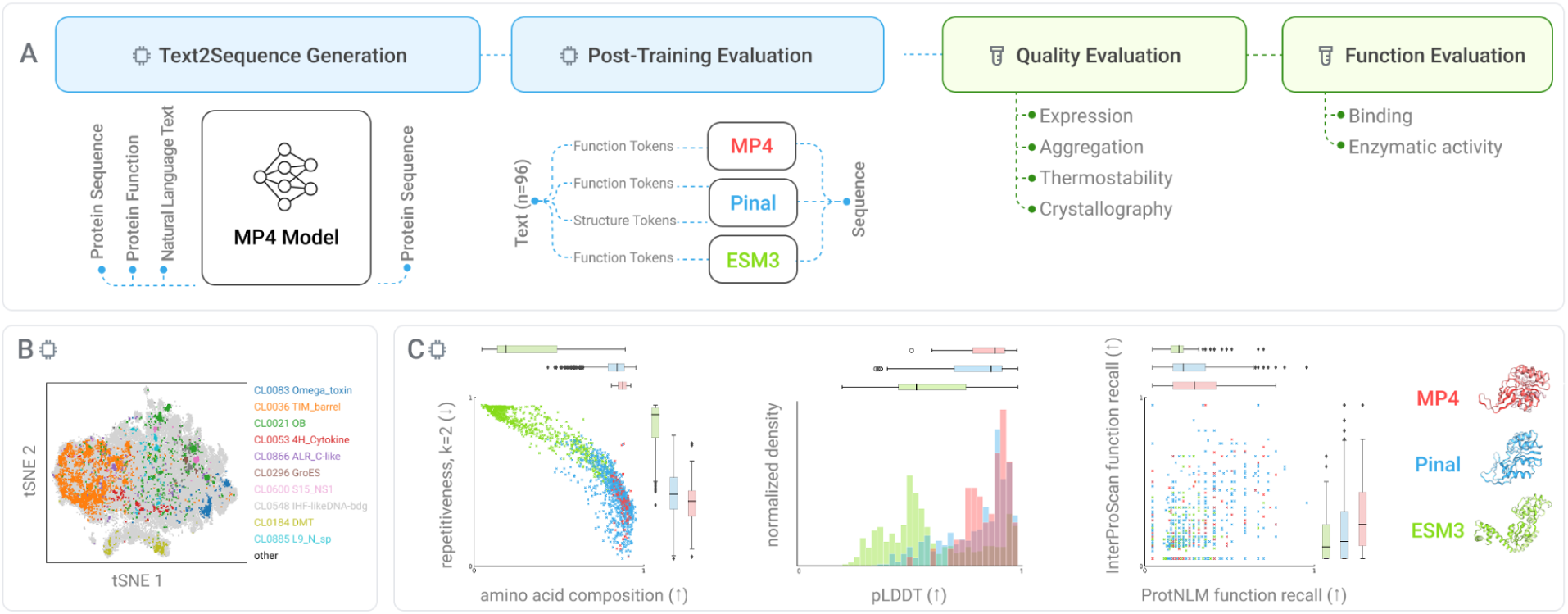
Overview of MP4 text-to-protein design and evaluation. A) Workflow: natural-language prompts are translated into sequences by MP4, followed by computational comparisons with Pinal and ESM3, and experimental assays for expression, stability, and function. B) t-SNE of MP4 embeddings highlighting distinct clustering of 10 Pfam clans. C) Quantitative comparison across 96 prompts. Left: sequence quality (amino acid composition vs. UniProt; repetitiveness k=2, lower is better). Center: predicted structure quality by ESMFold pLDDT. Right: functional recall (overlap between predicted functional keywords and those in the input prompt) using ProtNLM or InterProScan. Three representative structures from MP4, Pinal, and ESM3 are shown for the same prompt.

To assess how MP4 organizes protein information, we embedded natural proteins with MP4 and visualized the space with t-SNE(*18*). Distinct clustering by the Pfam clan can be found, indicating that MP4’s learned representation separates proteins by functional families (Fig. 1B)(*19*, *20*).

For benchmarking, we used ChatGPT to generate >1,000 prompts spanning enzymatic activities, binding partners, and subcellular localization, using vocabulary aligned to training ontologies (GO, InterPro, species descriptors, free-text)(*21–25*). From the resulting outputs, we selected 96 sequences for systematic comparison, prioritizing novelty (<50% identity to NR database) and diversity to avoid mode collapse (Fig. 2A)(*26*, *27*).

**Fig. 2.**
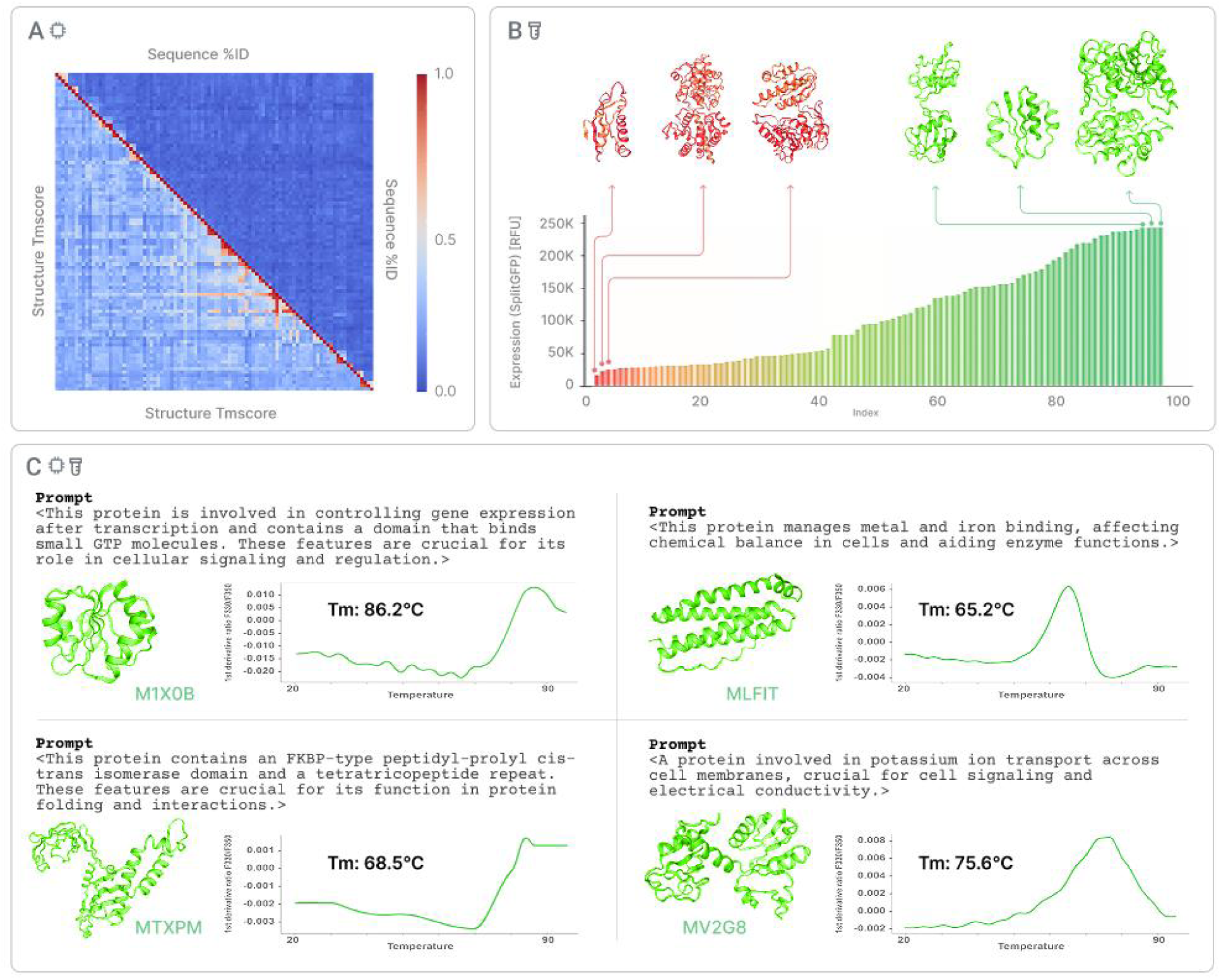
Experimental evaluation of 94 de novo designs. A) Heatmaps of pairwise similarity: sequence (upper right) and predicted-structure (lower left). B) Expression profile of all designs in a prokaryotic cell-free system, quantified by a split-GFP assay. C) Representative thermostability profiles measured by nanoDSF of four MP4-designed proteins.

To compare to ESM3, the 96 prompts were first transformed into InterPro classifications, which were then embedded, and input to ESM3(*9*). No sequence or structure templates are provided for a fair comparison to MP4. To compare to Pinal, the 96 prompts were directly input as text(*10*).

The prompts are used to create structural templates, which were used internally in Pinal to create the final sequences. Pinal therefore receives more starting information than either MP4 or ESM3, but as this step appears to be integral to Pinal, we did not disable the structure template step in this comparison.

We then evaluated sequence realism, predicted fold quality, and alignment to intended function (Fig. 1C). For the sequence, we examined amino acid composition (similarity to natural UniProt sequence distribution) and k=2 repetitiveness (lower is better)(*25*). MP4 outperformed both ESM3 and Pinal on both metrics. For structure quality, median ESMFold pLDDT was higher for MP4 designs than either ESM3 or Pinal(*28*). For alignment to the intended function, we standardized prompt terms and predicted annotations (from ProtNLM or InterProScan) to remove dashes, slashes, underscores, brackets, etc(*29*, *30*). Then we computed a recall-like score (shared terms divided by unique prompt terms). MP4 achieved higher scores than both ESM3 and Pinal across the panel. The last panel of Fig. 1C shows representative structures for a single prompt, visually illustrating the general observation that MP4 and Pinal seem to produce more similar results to each other than either does to ESM3.

Together, figure 1 shows that from text alone, MP4 generates sequences with native-like statistics, higher predicted structural confidence, and closer agreement between input prompt and predicted function than other models.

### High expression and thermostability across diverse de novo designs

We next asked whether MP4’s text-designed sequences yield experimentally tractable proteins. The 96-design panel was designed to span diverse sequence and predicted-structure space, as shown by pairwise similarity heatmaps (Fig. 2A). We cloned 94/96 constructs (two were unclonable due to higher-order repetitiveness) and expressed each in a prokaryotic cell-free system with a split-GFP assay(*31*). Most designs (79/94, 84%) produced detectable protein (Fig. 2B).

Next, we evaluated thermostability, a key property of designed proteins, by nanoDSF to determine the melting temperature (Tm)(*32*, *33*), the temperature at which 50% of the protein remains folded. Reliable melting curves were obtained for 17 designs (Table S7). Low intrinsic tryptophan limited nanoDSF signal for some designs, which could be overcome by prioritizing buried tryptophans during design. The proteins showed mean apparent Tm > 62 °C, with the most stable approaching 90 °C (Fig. 2C). To probe additional constructs lacking clear nanoDSF signals, we evaluated 10 designs by dynamic light scattering; each exhibited a single, uniform size distribution, consistent with well-behaved species in solution (Fig. S5)(*34*).

Together, these data show that MP4’s text-specified sequences frequently express and yield well-behaved, thermostable proteins in vitro across diverse predicted folds.

### Atomic-resolution structures validate MP4 designs and uncover novel topology

Crystallography—a gold standard in de novo protein design—provides definitive, atomic-level validation of text-designed sequences(*35*, *36*). We selected six highly expressed, thermostable MP4 designs for structure determination. Two yielded high-resolution X-ray structures (Fig. 3), establishing that text prompts alone can specify sequences that crystallize and resolve to atomic detail.

**Fig. 3.**
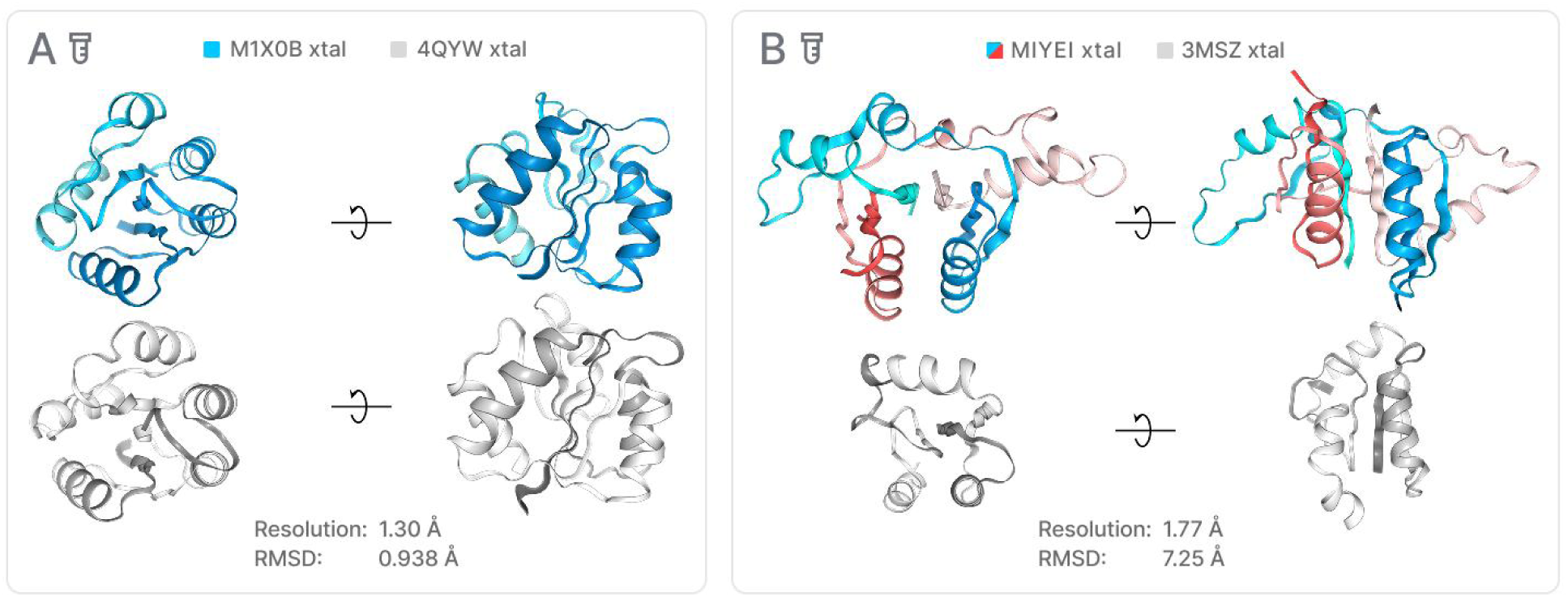
Crystal structures of MP4-designed proteins. A) M1X0B, solved at 1.30 Å, adopts a fold with close match to PDB ID 4QYW (RMSD 0.938 Å). B) MIYEI, solved at 1.77 Å, exhibits a novel fold with closest deposited structures 3MSZ and 3GPI at 7.25 and 5.53 Å RMSD, respectively.

M1X0B is a 120-amino acid monomeric, putative two-component response regulator solved at 1.30 Å. Its sequence is novel relative to the ClusteredNR database(ref) (54.7% identity to HPK26896.1), and the structure overlays closely with a known response-regulator-like fold (0.938 Å RMSD to PDB ID 4QYW)(*37*). By contrast, MIYEI demonstrates discovery beyond recall. This 77-amino acid, putative NAD(P)-bd_dom protein crystallized as a homodimer at 1.77 Å. It is novel in sequence (51.39% identity to WP_185270505.1 in ClusteredNR database) and structurally distinct, with its closest PDB matches (3MSZ, 3GPI) diverging by 7.25 Å and 5.53 Å RMSD, respectively—meaning MIYEI is a new fold.

While recent ML demonstrations (e.g., ESM3) highlight sequence-level novelty, crystal structures of M1X0B and MIYEI show that text-designed proteins are not only sequence-level novel, but also fold-level novel.

### MP4 designs functionally active ATPases

We next asked whether text-only prompts can yield biochemically active proteins. Many MP4 designs were predicted to bind and/or catalyze adenosine triphosphate (ATP). Given ATP’s central role in metabolism and biotechnology, we curated 107 ATP-associated designs from the >1,000-prompt set for detailed evaluation(*38*, *39*). In expression tests, 69/107 (64.5%) produced measurable protein in a prokaryotic cell-free system and 5/107 (4.7%) in *E. coli* (Fig. S6–S7).

To enable rapid binding measurements, we selected a focused set of 50 designs that combined high expression with sufficient intrinsic tryptophan signal and assessed ligand binding by nanoDSF in the absence of ligand and in the presence of 2 mM ATP or the non-hydrolyzable analog AMP-PNP (Fig. S8) (*40*, *41*). Binding of ligand is expected to stabilize the ATP-binding pocket, resulting in a measurable thermal shift.

We carried forward the six ATP binders, representing two mechanistic classes for detailed study (Fig. 4A). Specifically, three ATP-binding cassette (ABC) transporter nucleotide-binding domains (MB11S, M2RXT, MREGP) and three adenylate kinases (MS4BB, MBMLF, M17H6), as classified by ProtNLM. ESMFold models, coupled with DiffDock-predicted ATP pockets, and nearest natural analogs identified by Foldseek provided a structural basis for testing ATP activity (Fig. 4B)(*42*, *43*).

**Fig. 4.**
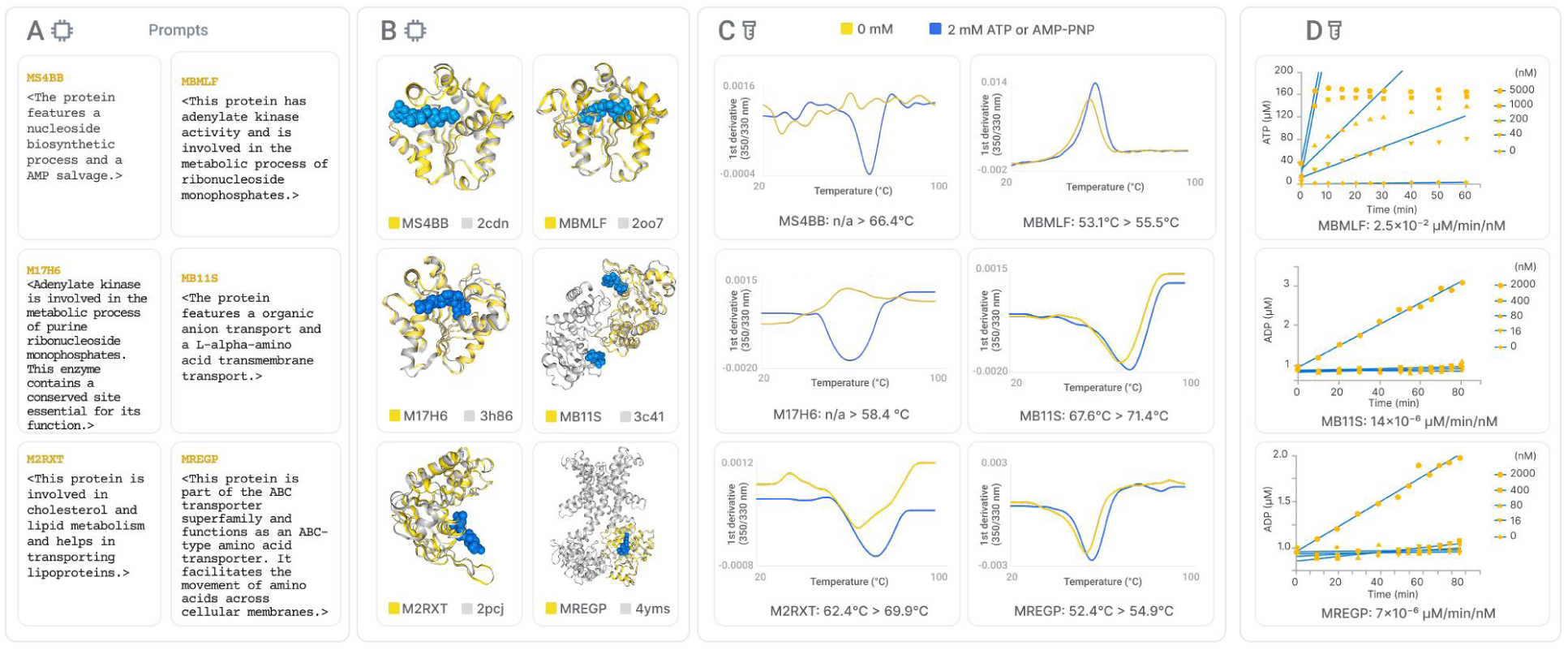
ATP binding and hydrolysis. A) Input prompts used for designs. B) Predicted structures (ESMFold; yellow), predicted ATP-binding pockets (DiffDock; blue), and nearest natural structural analogs (Foldseek; gray). C) Ligand binding assessed by nanoDSF as ΔTm upon addition of 2 mM ATP or AMP-PNP. D) ATPase activity for top candidates: ADP-Glo assay for adenylate kinase MBMLF and Kinase-Glo assay for ABC transporter domains MB11S and MREGP.

Ligand binding was detected as an increase in apparent Tm upon ligand addition, with differences of 2.4 °C in the case of MBMLF or up to 7.5 °C in the case of M2RXT (Fig. 4C). In two cases (MS4BB, M17H6), ligand addition converted previously featureless traces into well-defined melting transitions, consistent with binding-driven stabilization of the folded state.

We then tested catalysis in the top three candidates by expression level and thermal-shift confidence (Fig. 4D). Protein expression was scaled up in *E. coli,* and two different ATPase assays were used because it was determined that the Kinase-Glo assay (Promega) would be most straightforward for the ABC transporter domains designs (MB11S and MREGP) while the reversible nature of the adenylate kinase (MBMLF) activity was better suited for the ADP-Glo assay (Promega) to prevent interference of intermediate assay steps with final activity measurement(*44*, *45*). All three designs showed clear ATPase activity, confirming that MP4’s text-designed sequences hydrolyze ATP, not merely bind it.

These findings demonstrate that from natural-language prompts alone, MP4 produces proteins that bind ATP with measurable stabilization and catalyze ATP hydrolysis, meeting core biochemical criteria for function.

## Discussion

MP4 demonstrates that generalist, text-to-protein modeling can translate plain-language intent into sequences that score higher—simultaneously—on sequence realism, predicted fold quality, and alignment to requested functions than leading alternatives. In our head-to-head benchmark on 96 prompts, MP4 surpassed ESM3 and Pinal on amino acid composition and low repetitiveness, achieved higher median ESMFold pLDDT, and delivered closer agreement between prompt and predicted annotations. The overall design outputs from MP4 clustered more closely to Pinal than to ESM3, hinting at some shared understanding. Exploring the reasons underlying these patterns is an interesting future direction.

Importantly, this computational advantage translated into laboratory performance. Across a diverse set of de novo designs, most constructs expressed to detectable levels in a standardized cell-free system, many exhibited strong thermostability, and a subset crystallized to atomic resolution. Beyond folding, text-designed proteins measurably bound and hydrolyzed ATP in vitro, confirming their biochemical function. The six MP4-designed ATPases extends the small but growing canon of ML-designed enzymes—including pGAN59 (malate dehydrogenase; ProteinGAN), FastPETase (poly(ethylene terephthalate) hydrolysis; MutCompute), L056 (lysozyme; ProGen), and momi120_102 (serine hydrolase; PLACER = RFdiffusion+LigandMPNN) (*11*, *46–48*).

These results arrive as text-driven ML is reshaping how scientists specify and build. Shifting from template-based to text-based design can lower the barrier to protein engineering and widen the search space for discovery. Prompting remains an active challenge—and an opportunity—but MP4 shows that natural-language intent can yield proteins that express, crystallize, and catalyze(*49*, *50*). While early, MP4 points toward a new, low-barrier approach to protein design that invites more creativity - and more people - into the laboratory.

## Supporting information

mp4_pinal_esm3_comparison

## Acknowledgements

Experimental laboratory work was performed at Adaptyv Bio (programmable cloud lab), Tierra Biosciences, Proteros, and Arctoris Ltd. We thank Adaptyv Bio and Proteros for their crystallography expertise. We thank the team at Tierra Biosciences for technical expertise and project support. We thank Arctoris for using the Ulysses® platform, which generated high-quality, AI-ready data integral to this study on de novo proteins. We gratefully acknowledge the Arctoris team—particularly Eliot Osher, Devika Prabhakaran, Klem Simelis, Amit Sharma, and Rahul Yadav—for their expert execution of the experimental workflow and their invaluable scientific support. We thank Amédé Larabi, Kelvin Lau and Florence Pojer (Protein Production and Structure Characterization Core Facility, PTPSP, EPFL, Switzerland) for providing the resources for the protein expression and purification and for conducting the protein crystallization trials.We thank AMD Instinct Team for GPUs. AI-assisted technologies, just as ChatGPT was used for plotting and editing assistance.

## Author contributions

Conceptualization: KYW, KA, KA

Methodology: KYW, PK, IN, KA

Investigation: KYW, PK, IN, MS

Visualization: NB, MP, OM

Funding acquisition: KA

Project administration: KYW, TPR

Supervision: KYW

Writing – original draft: KYW, TPR, OM

Writing – review & editing: KYW, TPR, KA

## Competing Interests

The authors are employees of 310 AI.

## Data and materials availability

All data are available in the main text or the supplementary materials. Due to commercial confidentiality, the code is not publicly available. Researchers may contact 310 AI for access requests.

## Supplementary Materials

### Appendix A: MP4 Architecture

This appendix provides a high-level overview of the MP4 model architecture and its training regime. Due to proprietary considerations, specific implementation details—including internal layer configurations, precise hyperparameters, and optimization strategies—are not disclosed. The following sections outline the key design choices that enable MP4 to translate natural language prompts into functional, de novo protein sequences.

#### A.1 HIGH-LEVEL OVERVIEW

MP4 is a transformer-based model specifically designed for de novo protein design. It accepts natural language prompts that encode comprehensive protein information—such as fitness criteria, physical properties, source organism, and sequence-related properties—and generates protein sequences that meet the desired functional and structural constraints. The model emphasizes molecule programmability by integrating data from diverse sources and synchronizing multiple design objectives during training. This capability enables MP4 to deliver state-of-the-art performance in generating novel proteins that are both experimentally feasible and functionally robust.

**Fig. S1.**
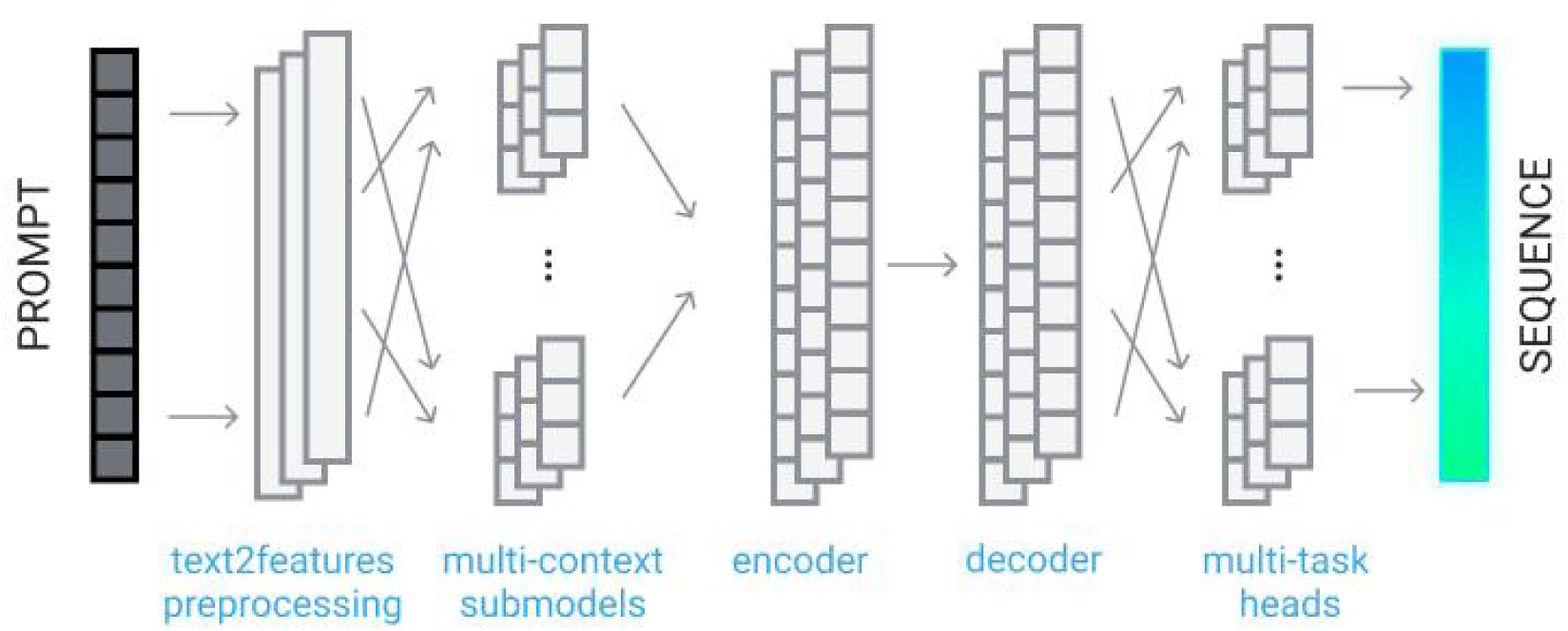
An overview of the MP4 architecture.

#### A.2 INPUT AND OUTPUT REPRESENTATIONS

**Input Representation:** The input to MP4 is a detailed textual prompt. These prompts include various descriptors such as:

- Fitness: Desired functional and performance metrics.
- Organism: Source organism or related biological context.
- Sequence Properties: Attributes that might include partial sequences, motifs, or structural hints.

Before being processed by the core model, these textual inputs pass through a text2feature preprocessing unit, which tokenizes the prompt and converts it into a structured feature representation. This conversion ensures that all pertinent information is captured and made accessible to subsequent layers.

**Output Representation:** The final output of MP4 is a protein sequence composed of amino acids. The sequence is generated in a way that reflects the functional and structural requirements encoded in the input prompt. By balancing multiple design tasks, MP4 ensures that the generated sequence is consistent with both the raw input and the underlying biochemical principles.

#### A.3 TRANSFORMER BACKBONE AND CONDITIONAL GENERATION

At the core of MP4 lies a multi-layer transformer architecture that has been adapted for the complexities of protein design. The architecture comprises several key components: Multi-Context Sub Models: These sub-models are designed to process different aspects of the input features. Each sub-model focuses on distinct information—derived from the prompt—ensuring that the model can attend to diverse functional, structural, and contextual cues.

- **Encoder:** After the multi-context sub models process the input, an encoder aggregates the information. It captures long-range dependencies and builds a rich, context-aware representation that summarizes the key aspects of the protein prompt.
- **Decoder:** The decoder translates the encoder’s latent representation into a form that is directly amenable to sequence generation. In this phase, the model transforms abstract feature representations into sequential data.
- **Multi-Task Head:** The output from the decoder is routed through a set of task-specific heads. MP4 is trained to perform 70 synchronized tasks, where each head is responsible for interpreting the decoder vector with respect to a particular design objective. These tasks include aspects such as structural fold determination, functional site prediction, and sequence novelty assessment. The collaborative output from these heads is then integrated to construct the final protein sequence.

Figure S1 schematically illustrates the end-to-end architecture of MP4, highlighting the sequential processing from prompt to final protein sequence.

#### A.4 TRAINING AND INFERENCE OVERVIEW

**Training Data and Regime:** MP4 has been trained on a highly diverse and extensive dataset, comprising over 3.2 billion datapoints (1.3 billion for sequences with length less than 300 and 2.3 billion for sequences with length less than 500) which were collected from various repositories, including UniProt. The training process involved processing 138K tokens and was carried out across 70 synchronized tasks, with each task emphasizing a distinct aspect of protein design—from structural features to functional specificity. Overall, the training was carried out with approximately 3,800 AMD-Instinct GPU-hours.

#### A.5 PROPRIETARY CONSIDERATIONS AND FUTURE DIRECTIONS

While this appendix outlines the overarching design and training strategy of MP4, many technical details remain confidential. In particular, specific modifications to the standard transformer framework, internal layer configurations, and fine-tuning strategies are proprietary. Future research will focus on:

- Refining the multi-context sub models to enhance the model’s sensitivity to nuanced protein features.
- Expanding the range of synchronized tasks to capture an even broader spectrum of protein functionalities.
- Exploring alternative decoding strategies to further improve sequence fidelity and novelty.
- Integrating all-atom protein structure generation to enable the direct production of detailed, three-dimensional protein models.

This continued evolution aims to push the boundaries of molecule programmability in protein design.

### Appendix B: Experimental Methods

#### B.1 CONSTRUCT DESIGN AND MOLECULAR CLONING

**Adaptyv:** Constructs were designed with C-terminal tags (GFP11 for split-GFP solubility and Twin-Strep for purification) and sourced from Twist Bioscience. The DNA constructs were subsequently assembled into the appropriate expression vector using the NEBuilder HiFi DNA Assembly Kit (New England Biolabs) according to the manufacturer’s protocol, with assembly reactions performed in 3 μL volumes.

**Tierra:** Construct design was performed using proprietary terminal DNA parts developed by Tierra Biosciences, which include solubility-enhancing and affinity purification tags. These tags were ligated to 310 AI-designed cDNAs using Golden Gate Assembly. The resulting assemblies were PCR amplified with proprietary primers to generate expressible DNA constructs encoding amino- (N-) and carboxy- (C-) terminal tagged protein constructs.

**Arctoris:** Genes encoding proteins of interest were designed with N-terminal 6xHis tag followed by TEV cleavage site and codon optimised for bacterial expression. Gene synthesis was performed at Genscript and subcloned into a pET-28a(+) vector (Genscript).

#### B.2 PROTEIN EXPRESSION AND PURIFICATION

**Adaptyv:** Protein expression was carried out in a prokaryotic cell-free system at Adaptyv Bio. Briefly, expression reactions were prepared in a total volume of 60 μL and incubated at 37 °C for 12 hours to ensure robust protein synthesis. Expression was detected by the fluorescent coexpression of the GFP11 tag and GFP1-10 marker as determined by a split-GFP solubility assay(*31*)). For protein purification, an affinity capture approach was employed using magnetic beads. Protein samples were mixed with the beads and incubated for 10 minutes at room temperature with gentle agitation to allow binding. Following binding, the beads were washed three times with a washing buffer composed of 1 M Tris-Cl, 1.5 M NaCl, and 10 mM EDTA (pH 8) to remove unbound proteins. The concentration and yield of the purified proteins were quantified using an affinity-based assay, with normalization performed via a Qubit fluorometer assay.

**Tierra:** Protein synthesis was carried out using expressible DNA fragments and Tierra Biosciences’ proprietary Enzyme and Biologics Optimized cell-free protein expression systems. Cell-free protein synthesis reactions were performed at 29 °C, followed by affinity purification using beads targeting the incorporated terminal tags. Proteins were subsequently cleaved from purification tags using a proprietary method and tag fragments were removed by centrifugation.

Final protein eluates were delivered in 50 μL of Tierra Biosciences storage buffer (100 mM Tris, 150 mM NaCl, 20% glycerol, pH 7.4).

Protein concentration was measured by UV/Vis spectroscopy using the Lunatic Spectrophotometer (Unchained Labs). Protein presence and purity were evaluated by SDS-PAGE. Gel electrophoresis samples were prepared using 4x Bolt LDS Buffer (Invitrogen) and 10x NuPAGE Sample Reducing Agent (Invitrogen). Samples were denatured at 70 °C for 10 min and separated on NuPAGE 4-12% Bis-Tris Protein Gels (Invitrogen) for 20 min at 200V using a Mini Gel Tank (Invitrogen). Gels were washed with water, stained for one hour with SimplyBlue SafeStain (Thermo Fisher Scientific), and destained in water. Gel images were captured using GelDoc Go (Bio-Rad) and Image Lab software (Bio-Rad) was used to assess sample purity.

**Arctoris:**

Plasmids were transformed into *E. coli* BL21 (DE3) STAR cells (Thermo Fisher Scientific) and grown in LB (0.5% glucose) (Melford) at 37 °C for 3 hr to OD600 = 0.6-0.8. The cells were induced with 0.5 mM IPTG (Sigma) and then grown further at 18 °C for 18 hr.

Cells were harvested by centrifugation at 4000 × g for 10 min, and lysed using Bug Buster buffer (Merck Millipore) containing benzonase (Sigma) and protease inhibitor tablets (Thermo Fisher Scientific). The lysate was clarified by centrifugation (15000 x g, 30 minutes, 4 °C) and subjected to His pulldown using His MultiTrap FF plate (Cytiva). Protein was eluted with 300 mM imidazole in PBS pH 7.3 and buffer exchanged with PBS using a PD MultiTrap G-25 plate (Cytiva).

For biochemical assays, cells were suspended (5 mL buffer/1 g cell pellet) in lysis buffer (50 mM Tris pH 7.8, 800 mM NaCl, 5% glycerol, 10 mM imidazole, 25 U/mL benzonase, protease inhibitor tablets, and 1 mM TCEP) and lysed using a cell disruption homogeniser (20 kPSI with one pass-through).

The lysate was clarified by centrifugation (45000 rpm, 45 minutes, 4 °C) and the His-tagged protein was captured on a 5 mL HisTrap FF crude column at a flow rate of 3 mL/min. The column was washed with 30 CV of wash buffer (50 mM Tris pH 7.8, 500 mM NaCl, 5% glycerol, 30 mM imidazole, 5 mM MgCl2, 1 mM TCEP) and the protein was eluted using a 0-100% linear gradient of elution buffer (50 mM Tris pH 7.8, 500 mM NaCl, 5% glycerol, 300 mM imidazole, 5 mM MgCl2, 1 mM TCEP) over 25 CV. Eluted fractions were analyzed by SDS-PAGE, and pure fractions were combined and concentrated to 5 mL using a 3 kDa MWCO ultra centrifugal filter. The protein was further purified by loading onto an S200 16/60 size exclusion column and eluting in size exclusion buffer (50 mM HEPES pH 7.5, 150 mM NaCl, 10 mM MgCl2, 5% glycerol). Eluted fractions were analyzed by SDS-PAGE, and pure fractions were combined and concentrated using a 3 kDa MWCO ultra centrifugal filter to an appropriate concentration. Final concentration was determined by measuring absorbance at 280 nm. Protein ID was confirmed by intact mass ESI MS on a Waters BioAccord LC-MS System. Concentrated protein was aliquoted, flash frozen in liquid nitrogen, and stored at -80 °C.

#### B.3 THERMOSTABILITY

**Adaptyv:** Protein thermostability was assessed using nano differential scanning fluorimetry (nanoDSF). Protein samples were diluted to a concentration of 100 μg/mL in assay buffer (20 mM sodium phosphate, 150 mM NaCl, pH 7.0), and 10 μL aliquots were pipetted into nanoDSF capillaries. The assay was performed with a temperature ramp of 1 °C per minute while monitoring intrinsic fluorescence, specifically by tracking the ratio of fluorescence intensities at 350 nm and 330 nm. Fluorescence changes were recorded continuously during the temperature increase, and the melting temperature (Tm) was determined as the inflection point on the fluorescence change curve. Tm values obtained for different constructs were compared to evaluate relative thermostability. Measurements reported were single measurements from a given sample.

**Arctoris:** Protein thermal stability in the presence and absence of AMP-PNP was assessed by nanoDSF on a Prometheus NT.48 (NanoTemper Technologies). Purified protein samples at 0.1–2 mg/mL in 10 mM HEPES pH 7.4, 150 mM NaCl, 10 mM MgCl_2_ were mixed with buffer or AMP-PNP (2 mM final concentration) (Sigma). 10 μL of protein solution was loaded into standard-grade nanoDSF capillaries (NanoTemper Technologies) and placed into the sample holder. Thermal unfolding was monitored by recording intrinsic fluorescence at 330 nm and 350 nm over a temperature range of 20 °C to 95 °C and a linear ramp of 1.5 °C/min. Fluorescence intensity ratio (F_350_/F_330_) was plotted as a function of temperature and the melting temperature (T_m_) was determined as the inflection point of the first derivative of F_350_/F_330_ using PR.ThermControl software (v2.3.1, NanoTemper Technologies).

#### B.4 DYNAMIC LIGHT SCATTERING

**Adaptyv:** Dynamic light scattering (DLS) was used to evaluate the hydrodynamic radius and aggregation state of the expressed proteins. Prior to measurement, purified protein samples were allowed to equilibrate to room temperature. Approximately 20 μL of each sample was loaded into a disposable cuvette, and measurements were performed at 25 °C using the Unchained Labs Uncle instrument. A series of at least 10 runs per sample was acquired to ensure statistical reliability. The size distribution data were analyzed to determine the hydrodynamic radius and to assess the presence of protein aggregates. Measurements reported were single measurements from a given sample.

#### B.5 ADENYLATE KINASE ACTIVITY ASSAY

**Arctoris:** Adenylate kinase activity was measured using the Kinase-Glo assay kit from Promega. Reagents were dispensed into a 384-well low volume, white plate (Greiner, 784075) using the Dragonfly liquid handler (SPT Labtech). Buffer consisting of 50 mM HEPES (pH 7.5), 10 mM MgCl2, 1 mM DTT, 0.5% Glycerol, 0.02% BSA, 0.003% Tween-20, and 0.5 mM EGTA was dispensed into the plate with volumes ranging from 0.625 µL to 10 µL for dilution of constructs and mitigation of evaporation. Corresponding volumes of Construct 95 (49 µM) was added to those wells to create a 5-fold dilution series of final enzyme concentrations (5 µM, 1 µM, 200 nM, 40 nM) to a volume of 5 µL at which point the plate was centrifuged at 200 g for 60 sec prior to addition of another 5 µL of 1 mM ADP (final concentration 500 µM) to initiate the reactions. ADP additions were carried out at specified timepoints to provide data for 0, 5, 10, 15, 20, 25, 30, 40, 50, 60 minute reaction durations. Upon completion of the time course, 5 µL of Kinase Glo reagent was added to each well following a 60 sec 200 g centrifugation and incubated for 10 min. The plate was measured using a PHERAstar FSX plate reader (BMG Labtech). An ATP standard curve generated during the onboarding process was then used to convert the relative luminescent unit (RLU) signals to final ATP concentration in each well.

#### B.6 ATPASE ACTIVITY ASSAY

Arctoris: ATPase activity was measured using the ADP-Glo assay kit from Promega. Reagents were dispensed into a 384-well low volume, white plate (Greiner, 784075) using the Dragonfly liquid handler (SPT Labtech). Buffer consisting of 50 mM HEPES (pH 7.5), 10 mM MgCl2, 1 mM DTT, 0.5% Glycerol, 0.02% BSA, 0.003% Tween-20, and 0.5 mM EGTA was dispensed into the plate with volumes ranging from 0.625 µL to 10 µL for dilution of constructs and mitigation of evaporation. Corresponding volumes of Constructs 15 (64 µM) and 64 (18 µM) were added to those wells to create a 5-fold dilution series of final enzyme concentrations (5 µM, 1 µM, 200 nM, 40 nM) to a volume of 5 µL at which point the plate was centrifuged at 200 g for 60 sec prior to addition of another 5 µL of 400 µM ATP (final concentration 200 µM) to initiate the reactions. ATP additions were carried out at specified timepoints to provide data for 0, 10, 20, 30, 40, 50, 55, 60, 65, 70, 75, and 80 minute reaction durations. Upon completion of the time course, 5 µL of Reagent 1 from the ADP-Glo assay kit was added to each well following a 60 sec 200 g centrifugation and incubated for 60 min. 10 µL of Reagent 2 from the ADP-Glo assay kit was then added to all wells, centrifuged at 200 g for 60 sec, and incubated for 60 min before measuring using a PHERAstar FSX plate reader (BMG Labtech). An ADP standard curve generated during the onboarding process was used to convert the relative luminescent unit (RLU) signals to final ADP concentration in each well.

#### B.7 X-RAY CRYSTALLOGRAPHY

**Adaptyv:**

*Protein expression and purification:* The M1X0B(3best9) expression plasmid carrying a C’ terminal 8xHistidine tag was cloned into the bacterial expression vector pET-28a(+). The coding sequence was optimized for *Escherichia coli* (*E. coli*) expression. The plasmid was transformed into BL21(DE3) competent cells. Protein expression was induced with 0.4 mM isopropyl b-D-1-thiogalactopyranoside (IPTG) when cells reached an optical density (OD600) of 0.6, with a subsequent overnight growth at 18 °C. The cell pellet was resuspended in lysis buffer (500 mM NaCl, 20 mM HEPES pH 7.4, 5% glycerol, supplemented with cOmplete™, EDTA-free Protease Inhibitor Cocktail, Roche) and was lysed by sonication. After centrifugation at 20,000 x g for 60 minutes at 4°C, the supernatant was filtered and applied to a HisTrap excel column (Cytiva).

After elution of recombinant 3best9 protein with a continuous gradient over 25 column volumes of elution buffer (500 mM NaCl, 20 mM HEPES pH 7.4, 5% glycerol, 500 mM imidazole), pure fractions were dialyzed overnight with 3 kDa cutoff membrane against 5 L of final buffer (150 mM NaCl, 20 mM HEPES pH 7.4). Dialyzed protein was concentrated, and additionally purified by size-exclusion chromatography (HiLoad 16/600 Superdex 200 pg, Cytiva) in the final buffer. Pooled fractions were concentrated to the required concentration using a MWCO concentrator (Millipore) with a cutoff of 3 kDa.

*Protein crystallization and structure determination:* The 3best9 protein was crystallized at a concentration of 12.5 mg/ml using the sitting-drop vapor diffusion method at 18 °C. Crystallization drops were prepared in a 1:1 ratio with reservoir solution containing 0.2 M potassium sodium tartrate tetrahydrate, 2.0 M ammonium sulfate and 0.1 M sodium citrate, pH 5.6 (SG1 Screen HT-96, Molecular Dimensions). Crystals were cryoprotected in 25% glycerol and flash-frozen in liquid nitrogen. X-ray diffraction data were collected at -173.15 °C (100 K) on the ID23-1 beamline at The European Synchrotron (ESRF, Grenoble, France). Phases were obtained by molecular replacement using PHASER(*51*) with the 3best9 AlphaFold 3(*52*) predicted model as template. Model building and refinement was completed using COOT(*53*) and REFMAC(*54*). The quality of refined models was assessed using MolProbity(*55*).

**Table S1.** Data Collection Statistics for M1X0B.

|  | M1X0B (3best9) |
| --- | --- |
| Wavelength (Å) | 14.000 KeV (0.8856) |
| Space group | P212121 |
| Cell dimensions |  |
| a, b, c (Å) | 45.11, 45.22, 59.45 |
| $\alpha, \beta, \gamma$ (°) | 90, 90, 90 |
| Resolution (Å) | 1.30 (1.36 – 1.30) |
| Unique reflections | 30590 |
| $R_{\text{meas}}$ | 6.7% (317.4%) |
| $I / \sigma I$ | 16.76 (0.97) |
| Completeness (%) | 100% (100%) |
| Redundancy | 13.0 |
| CC1/2(%) | 99.9 (35.7) |

**Table S2.**
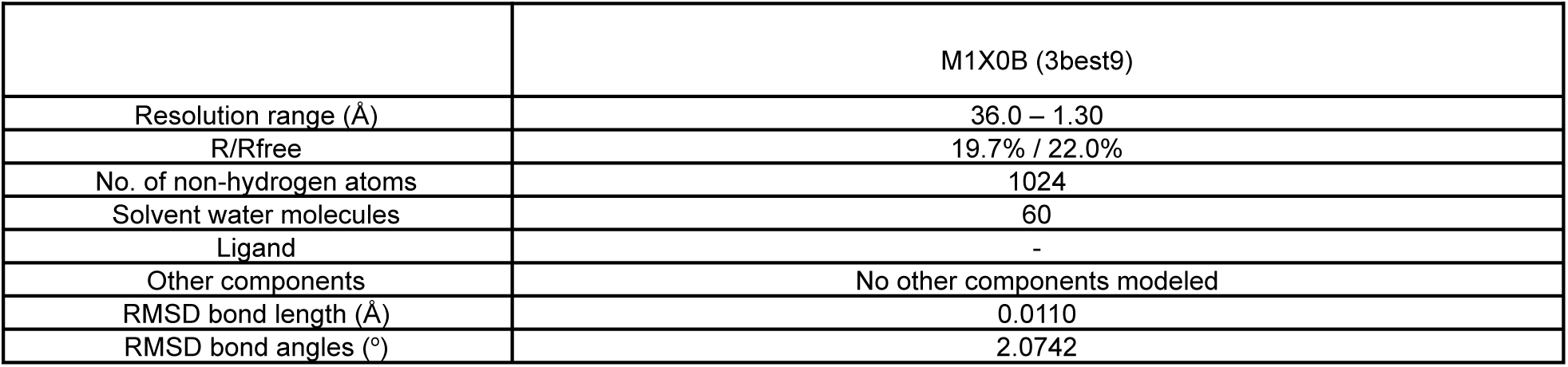
Refinement Statistics for M1X0B.

**Proteros:**

*Protein Production:* Suitable constructs for MIYEI expression have been established by Proteros. Expression of MIYEI and a purification protocol has been established. Homogeneous protein was produced in preparative amounts. The protein was purified comprising affinity and gel filtration chromatography steps. This procedure yielded homogenous protein with a purity greater than 95% as judged from Coomassie stained SDS-PAGE.

*Crystallization:* The purified protein was used in crystallization trials employing both, a standard screen with approximately 1200 different conditions, as well as crystallization conditions identified using literature data. Conditions initially obtained have been optimised using standard strategies, systematically varying parameters critically influencing crystallization, such as temperature, protein concentration, drop ratio, and others. These conditions were also refined by systematically varying pH or precipitant concentrations. The crystal has been harvested from a drop containing PEG20000, PEG550MME, bizine, sodium formate, ammonium acetate, trisodium citrate, sodium potassium L-tartrate, sodium oxamate, HEPES-NaOH, sodium chloride, DTT, DMSO, Trizma base at pH 8.5.

*Data Collection and Processing:* A cryo-protocol was established using Proteros Standard Protocols. Crystals have been flash-frozen and measured at a temperature of 100 K. The X-ray diffraction data have been collected from crystals of MIYEI at the European Synchrotron Radiation Facility (ESRF, Grenoble, France) using cryogenic conditions. The crystals belong to space group C 2. Data were processed using the programs autoPROC, XDS and autoPROC, AIMLESS.

The quaternary structure of artificial protein MIYEI consists of a homo dimeric protein complex with a six-stranded-sheet surrounded by six short helices. The C-terminal -strand of one monomer is inserted into the -sheet structure of the other monomer, allowing for the formation of a stable dimer. There is a homo dimer in the asymmetric unit with basically the same overall conformation. The model comprises the residues Ser2 to Gly79 (Table S4).

**Table S3.** Completeness of Model MIYEI.

|  |
| --- |
| Amino acid residues defined by electron density |
| A: 3-79 |
| B: 2-79 |
| Except for amino acid residues<sup>1</sup> |
| A: - |
| B: - |
| Except for side chains<sup>2</sup> |
| A: 5, 14, 38, 49 |
| B: 5, 21, 29, 45, 48-49, 61, 68 |

<sup>1</sup> Residues not included in the model
<sup>2</sup> Occupancy set to 0.00 in the model

**Table S4.**
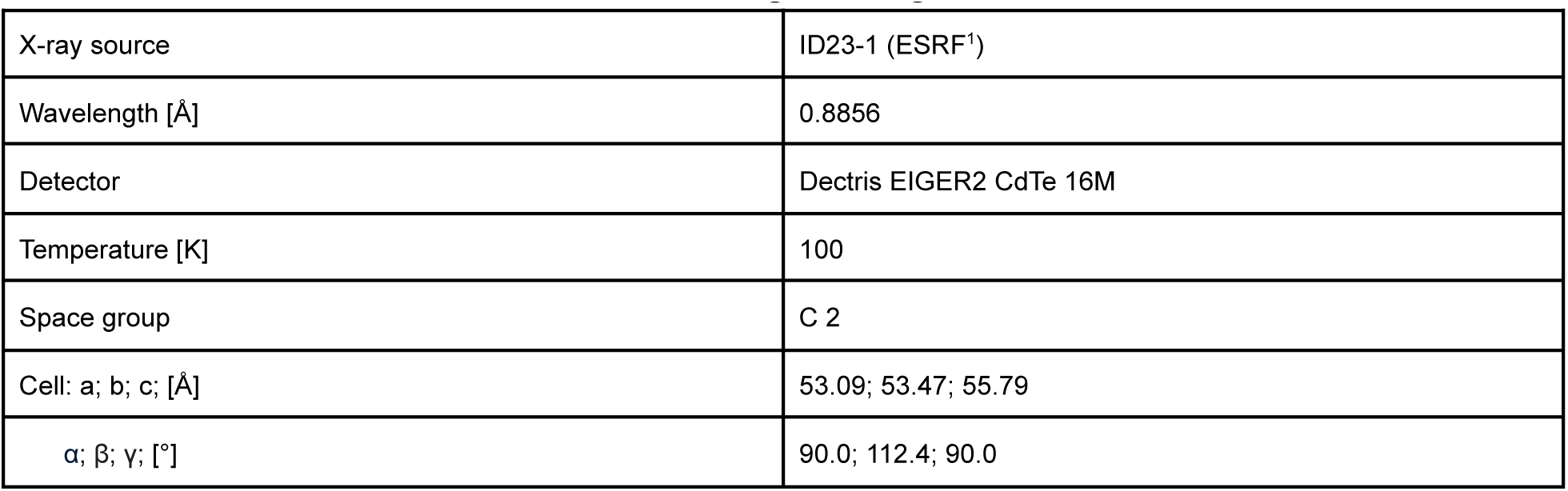
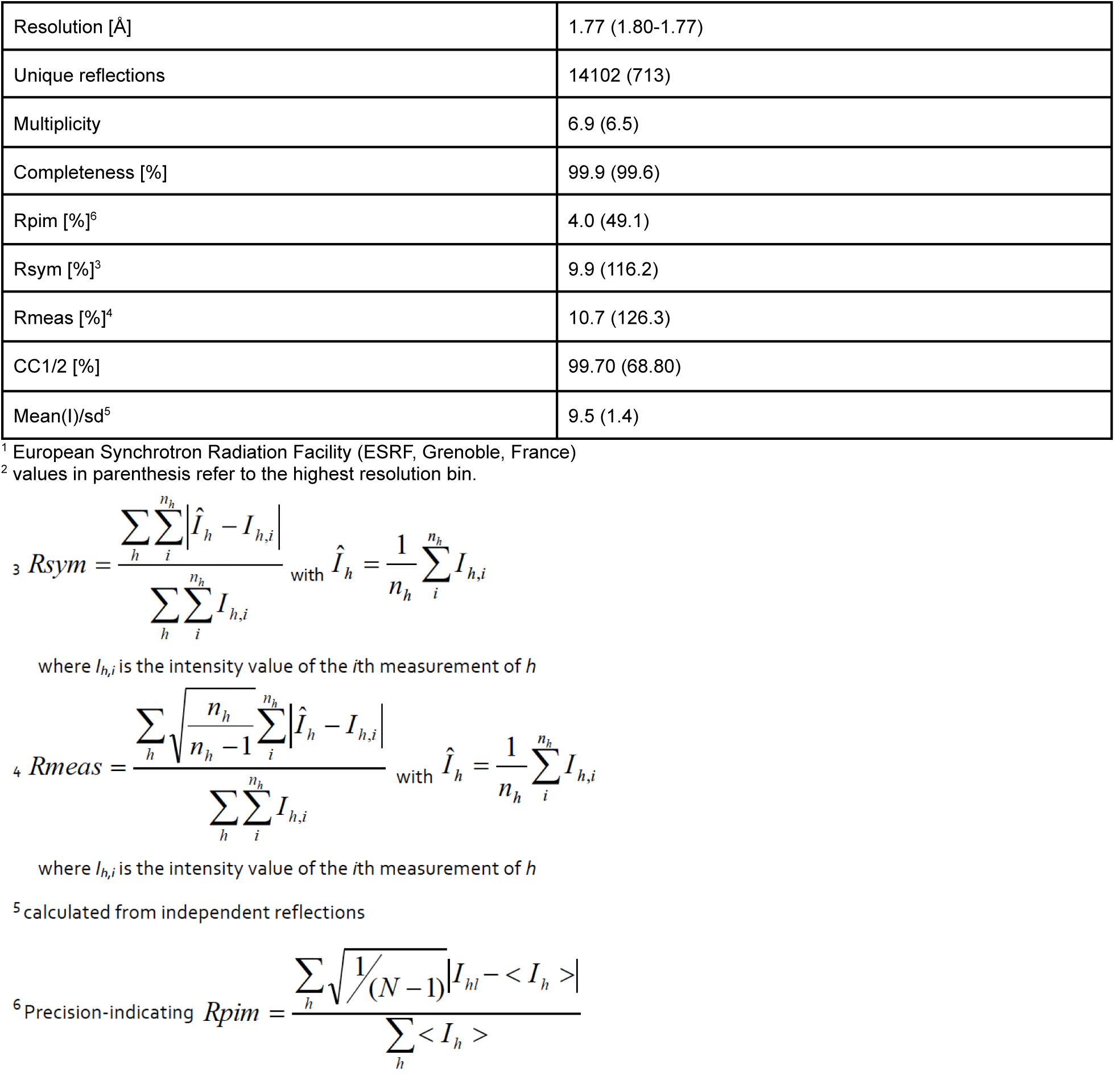
Data collection and processing statistics for MIYEI.

*Structure Modelling and Refinement:* The phase information necessary to determine and analyze the structure was obtained by molecular replacement. Structure model provided by the customer was used as a search model to solve the structure by Molecular Replacement method.

Subsequent model building and refinement was performed according to standard protocols with COOT and the software package CCP4, respectively. For the calculation of the free R-factor, a measure to cross-validate the correctness of the final model, about 5.0% of measured reflections were excluded from the refinement procedure (see Table S5).

Automatically generated local NCS restraints have been applied (keyword “ncsr local” of newer REFMAC5 versions).

The water model was built with the “Find waters"-algorithm of COOT by putting water molecules in peaks of the Fo-Fc map contoured at 3.0, followed by refine-ment with REFMAC5 and checking all waters with the validation tool of COOT. The criteria for the list of suspicious waters were: B-factor greater 80 Å, 2Fo-Fc map less than 1.2, distance to closest contact less than 2.3 Å or more than 3.5 Å. The suspicious water molecules and those in the ligand binding site (distance to ligand less than 10 Å) were checked manually.

The Ramachandran Plot of the final model calculated with MolProbity shows 98.01 % of all residues in the favored region and 1.99 % in the allowed region (Fig-ure 3). There are no outliers in the Ramachandran plot (Table S4). Statistics of the final structure and the refinement process are listed in Table S5.

**Table S5.**
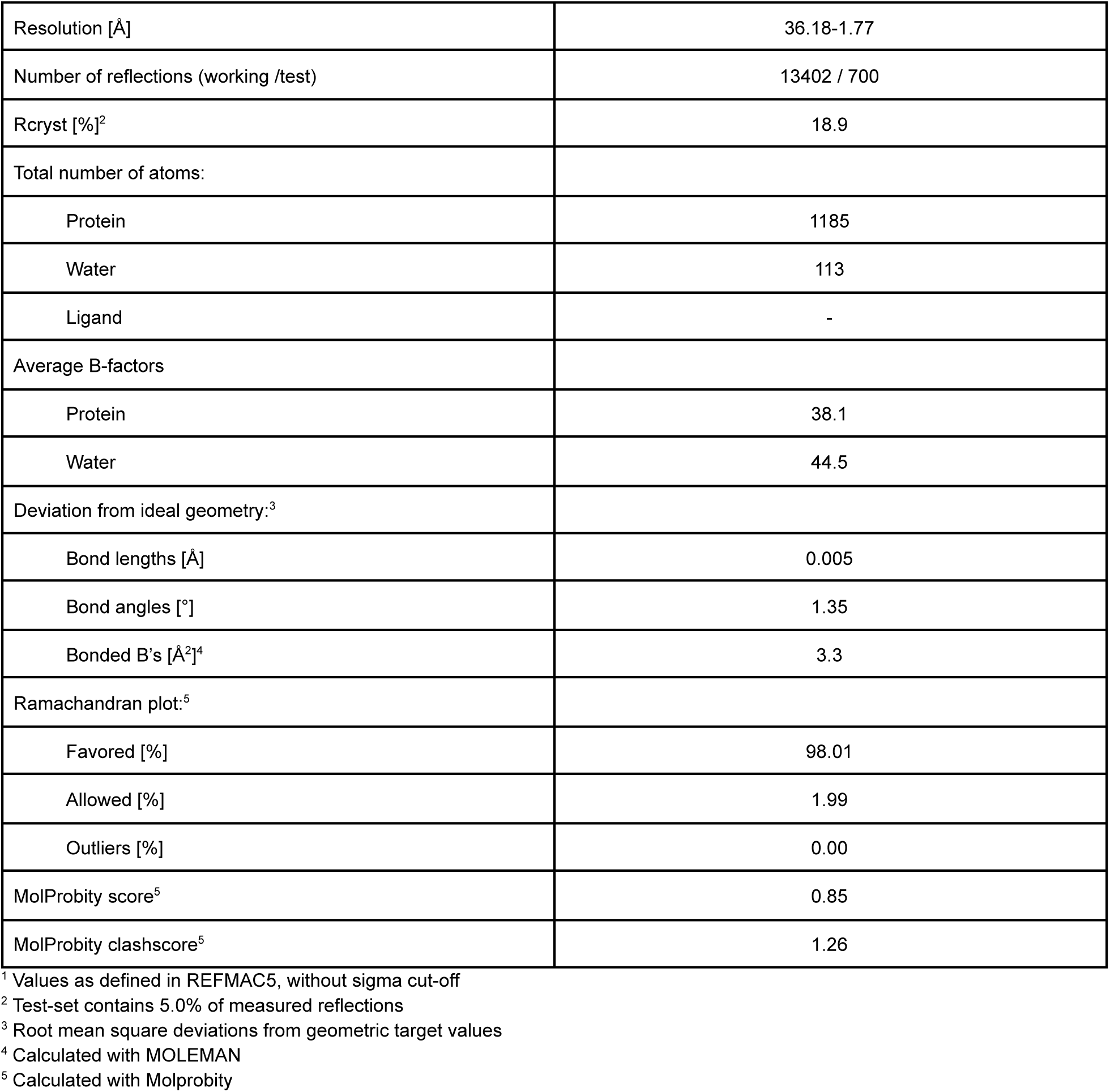
Refinement statistics for MIYEI^1^.

### Appendix C: Supplemental Data

**Fig. S2.**
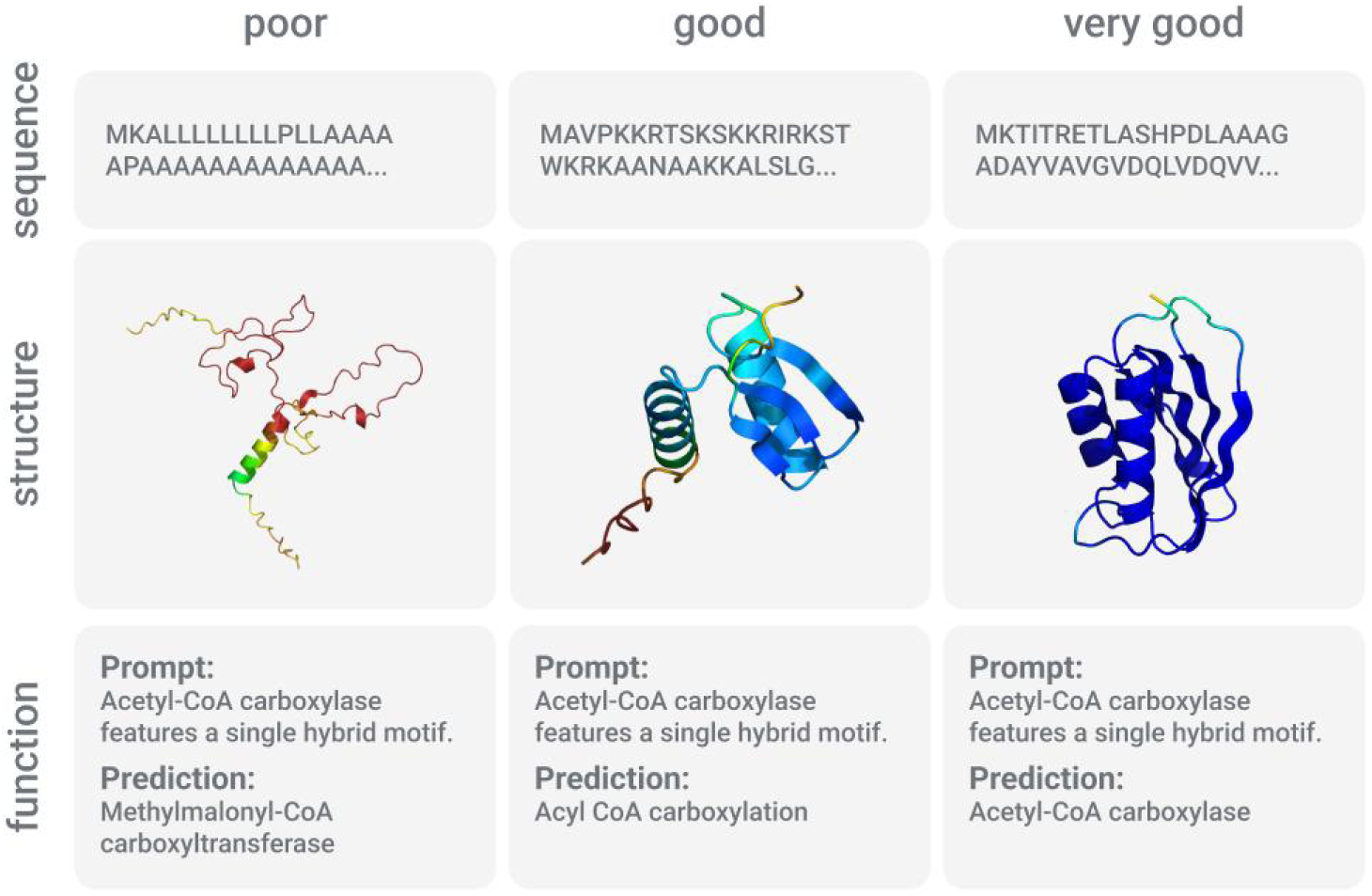
Examples of poor, good, or very good quality sequences, structures, and functions.

**Fig. S3.**
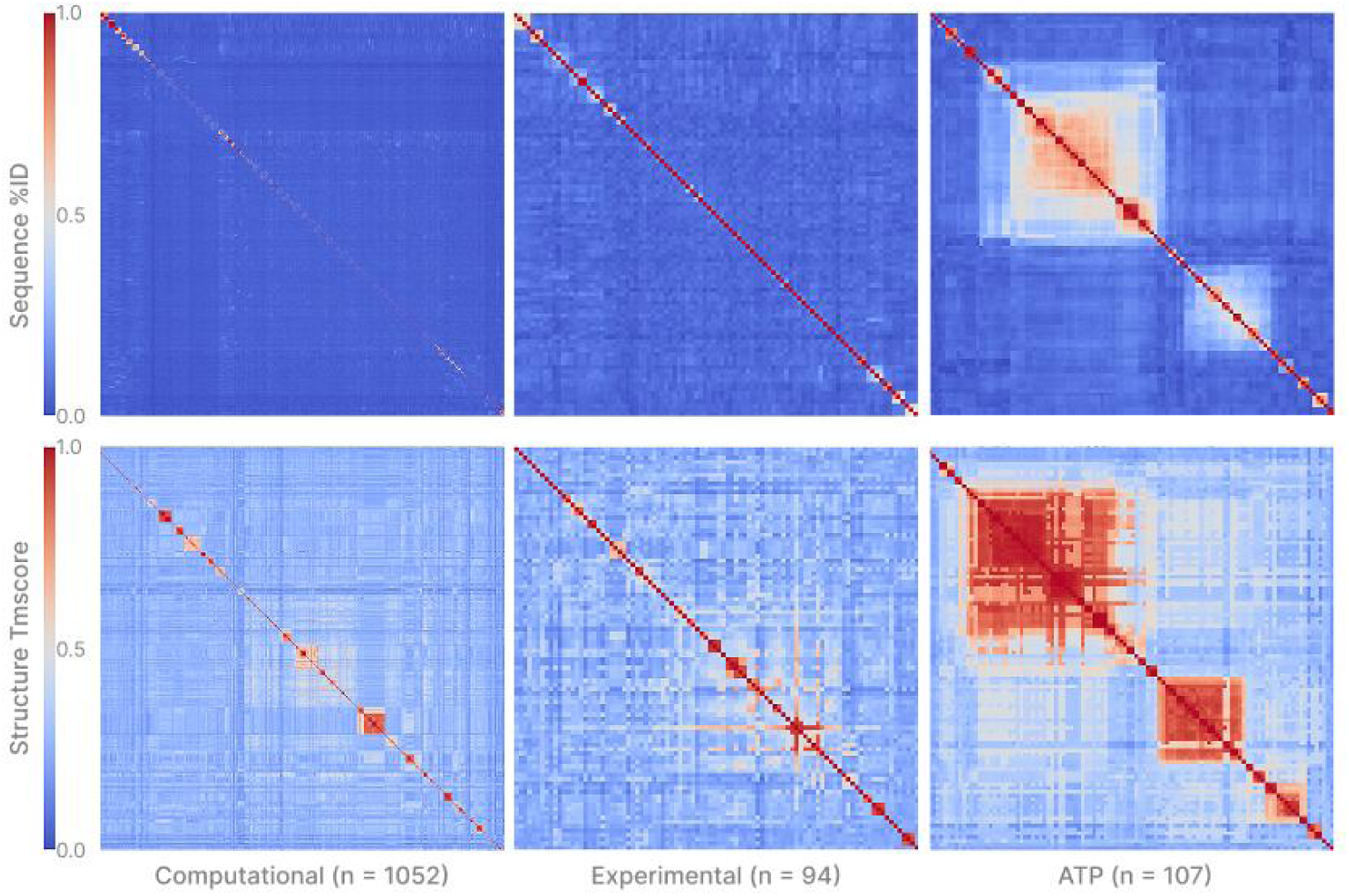
Heatmaps showing sequence (top) and structural (bottom) similarity of MP4 designed proteins compared to others in the full computational set (left), experimental validated diversity set (middle), and experimentally validated ATP activity focused set (right).

**Fig. S4.**
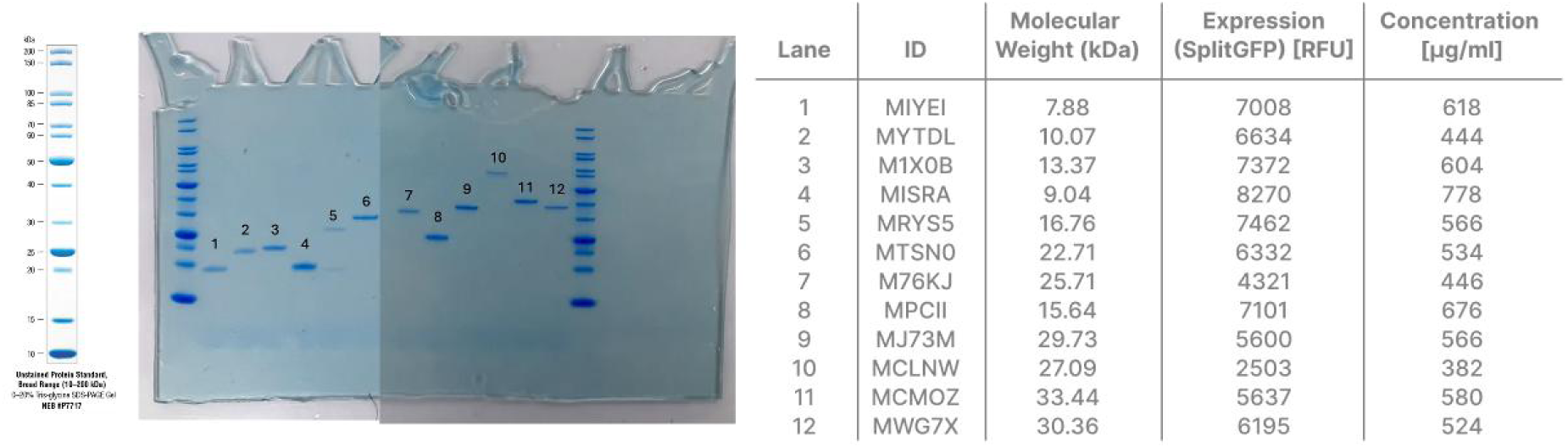
SDS-PAGE gel characterization of select proteins of the diversity set.

**Fig. S5.**
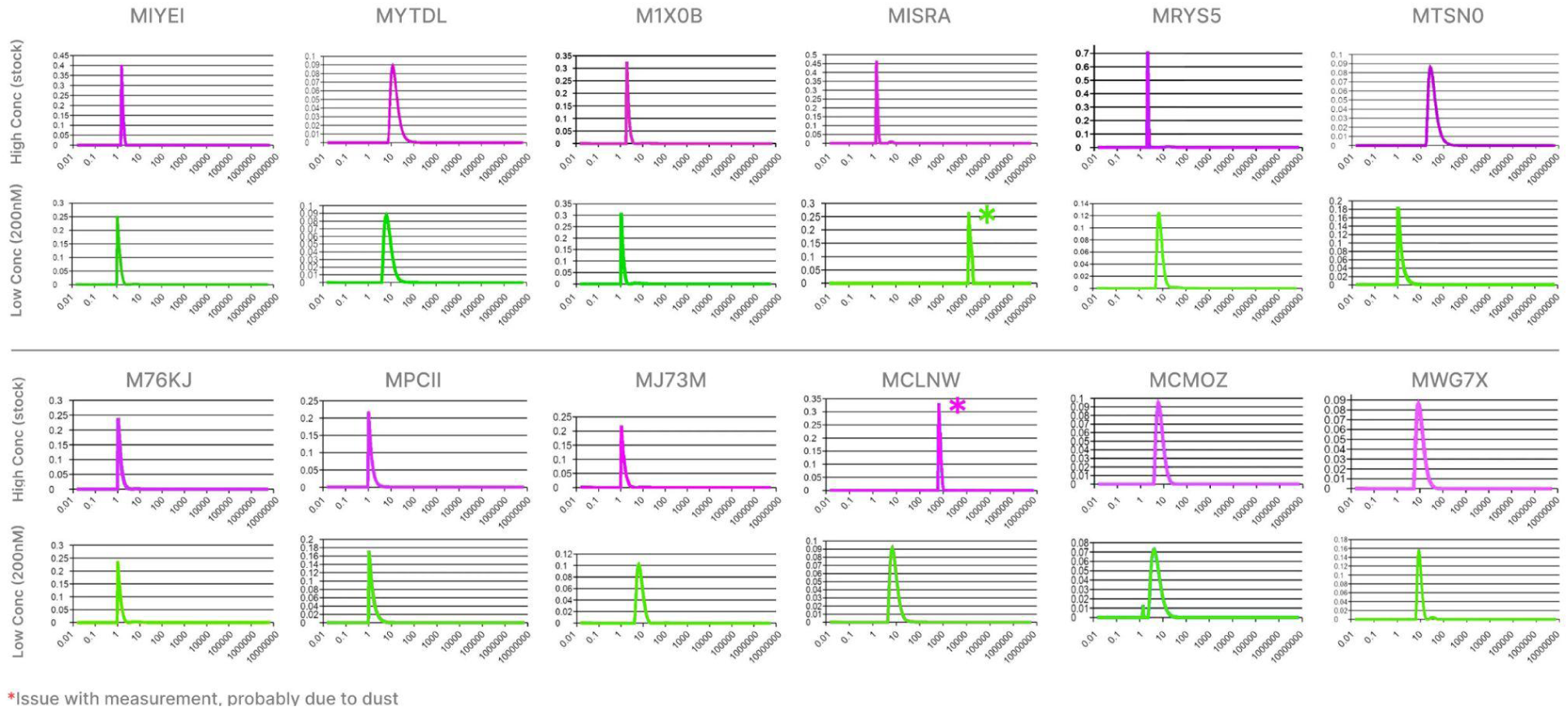
Dynamic light scattering (DLS) characterization of select proteins of the diversity set.

**Fig. S6.**
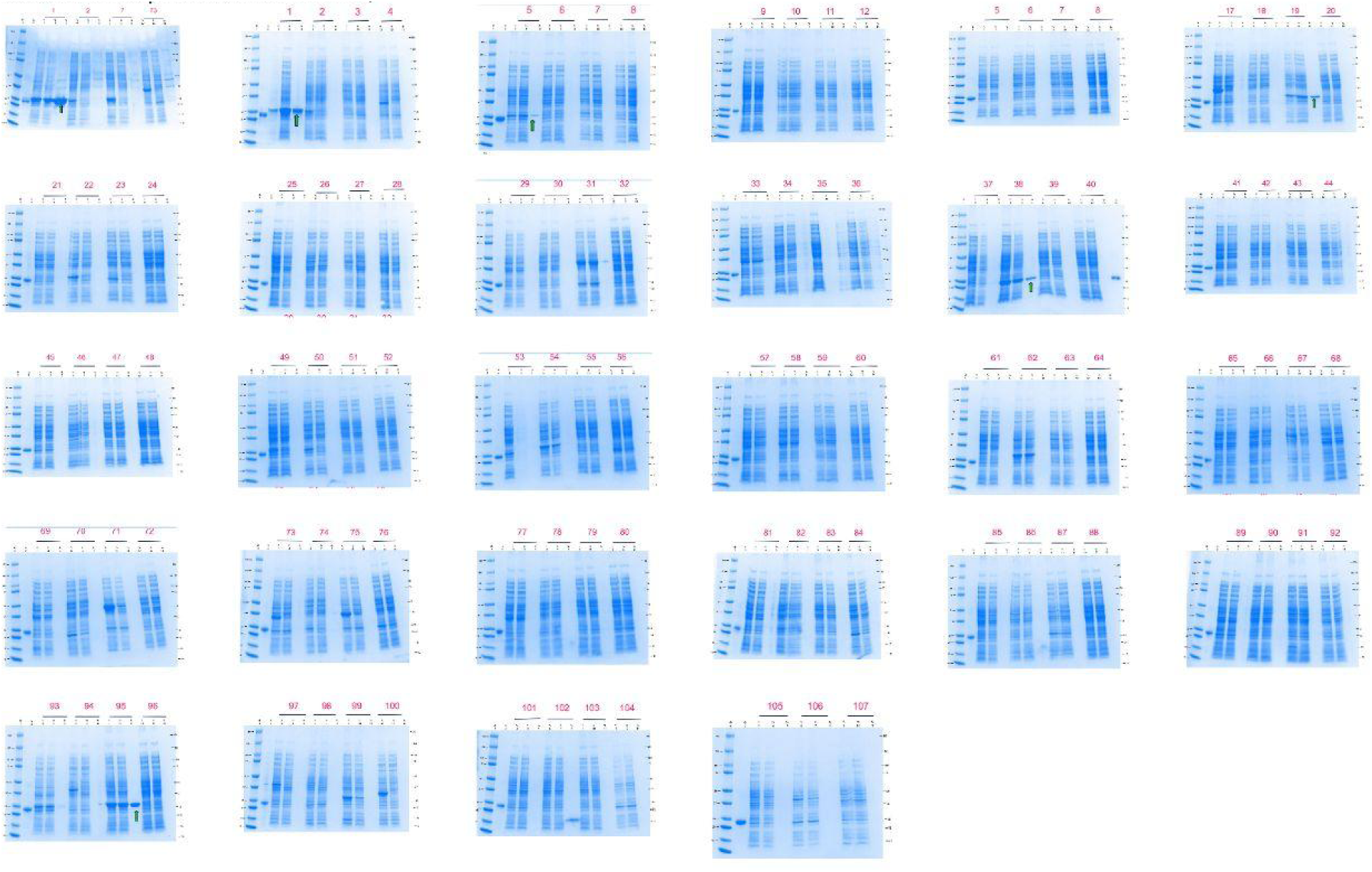
SDS-PAGE gel characterization of bacterial expression of the ATP activity set.

**Fig. S7.**
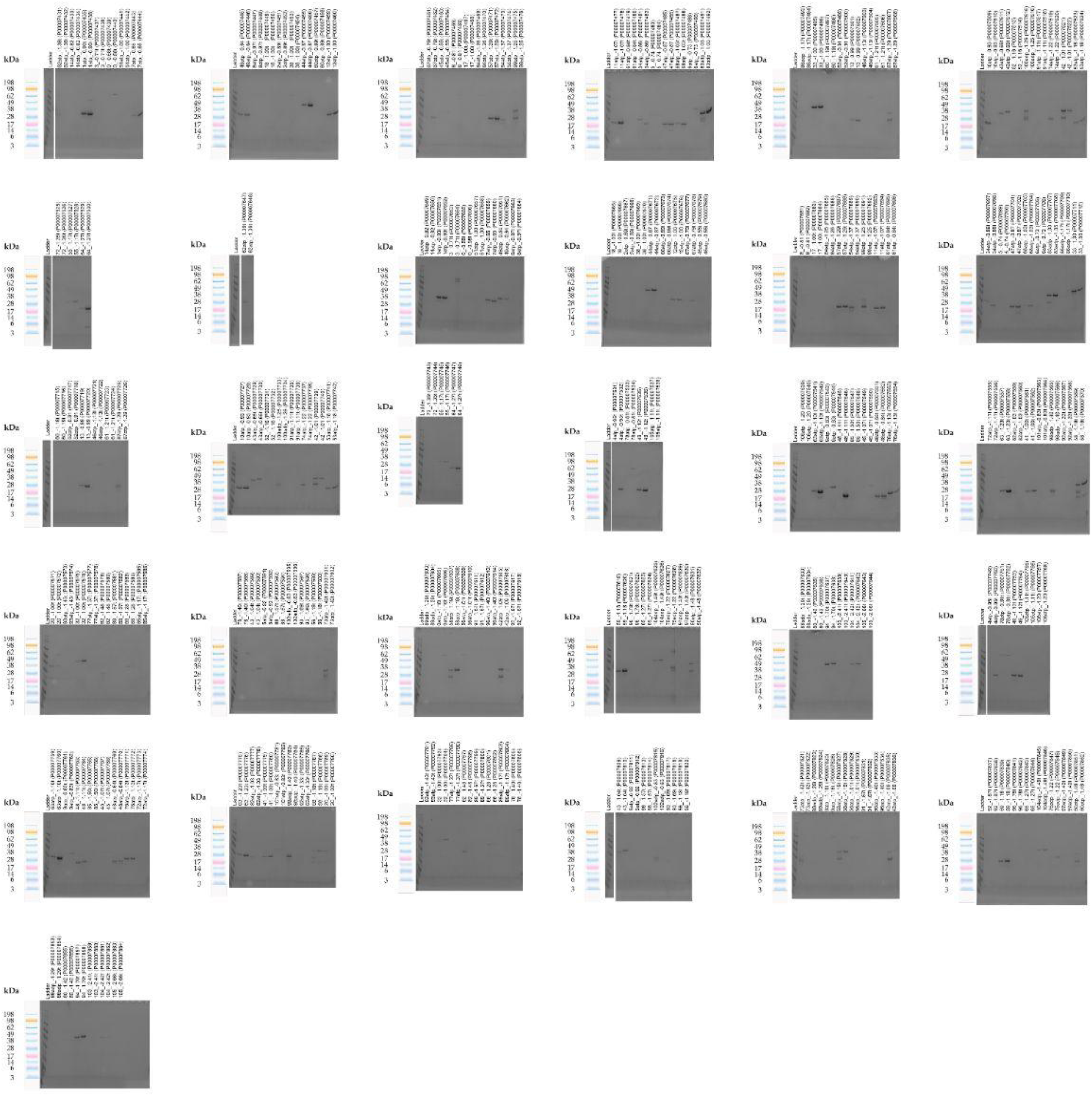
SDS-PAGE gel characterization of cell-free expression of the ATP activity set.

**Fig. S8.**
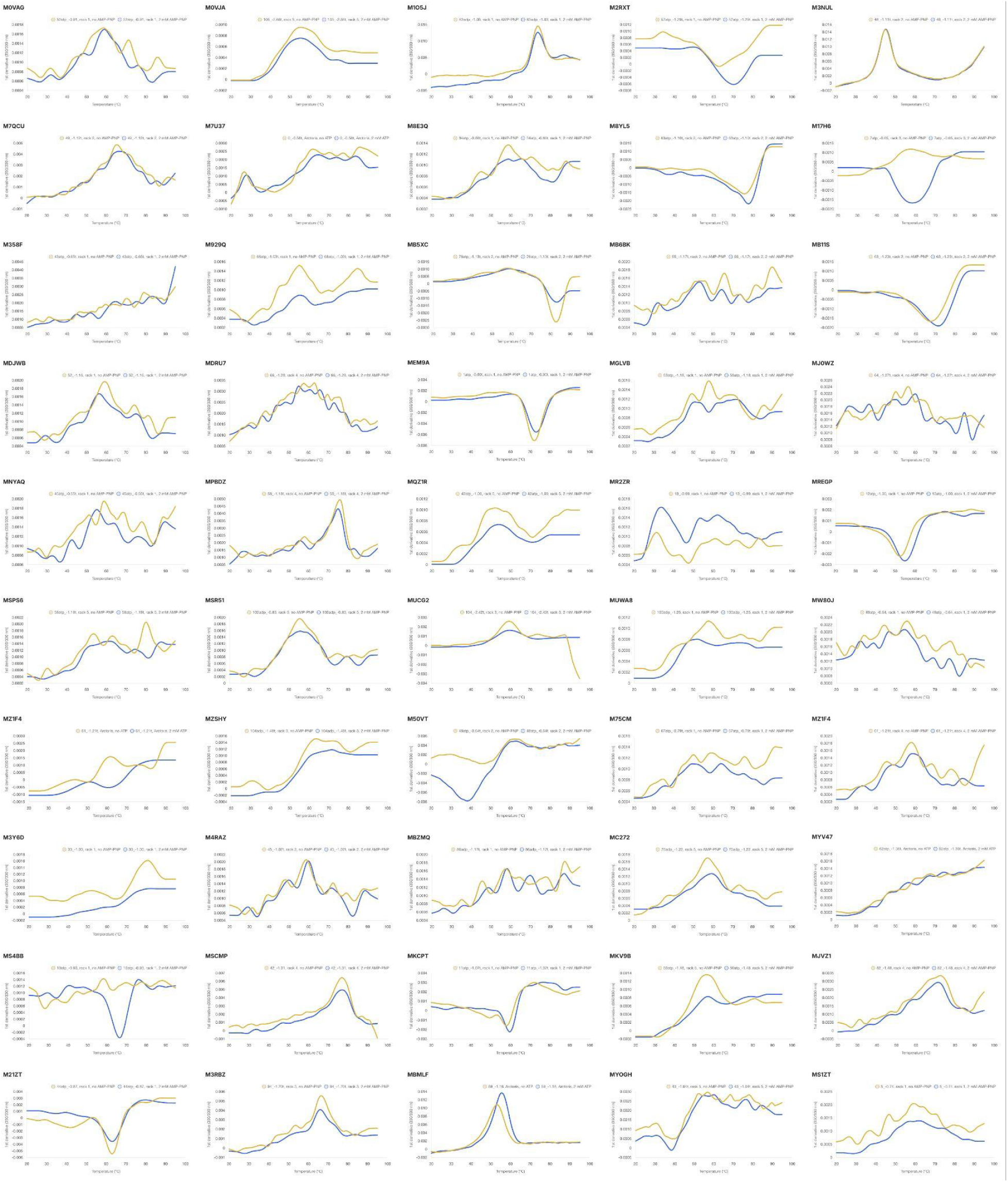
**Melting temperature by nanoDSF with 0 mM or 2 mM ATP or AMP-PNP of proteins in the ATP activity set.**

**Fig. S9:**
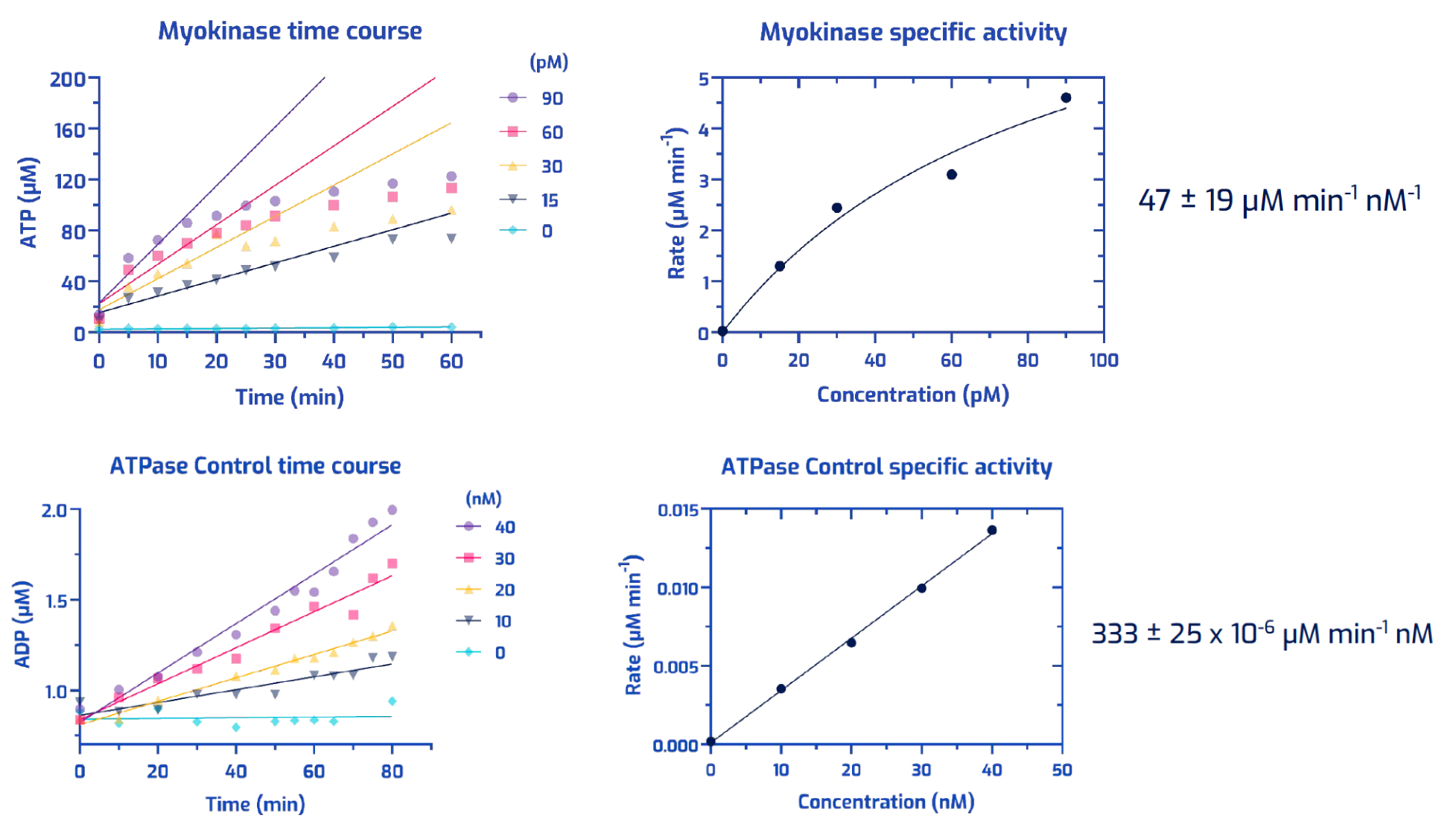
**Control protein rabbit muscle myokinase for adenylate kinase assay (top) and control protein MBPR1B for ADP-Glo assay (bottom)**

**Table S6.** Sample naming conversion table.

| Repo ID | Sample Name | Sample # | Repo ID | Sample Name | Sample # | Repo ID | Sample Name | Sample # |
| --- | --- | --- | --- | --- | --- | --- | --- | --- |
| M0VAG | 52atp_-0.91 | 35 | MAOYP | 72atp_-1.17t | 63 | MOFBD | 88_-1.57t | 82 |
| M0VJA | 105_-2.66t | 107 | MB11S | 63_-1.23t | 64 | MPBDZ | 58_-1.18t | 70 |
| M17H6 | 7atp_-0.65 | 7 | MB5XC | 76atp_-1.13t | 62 | MQU7R | 85_-1.50t | 59 |
| M17PF | 78_-1.40t | 79 | MB6BK | 55_-1.17t | 49 | MQZ1R | 42atp_-1.09 | 93 |
| M1CE2 | 61atp_-0.84t | 25 | MBGJG | 8atp_-0.97t | 9 | MR2ZR | 13_-0.99 | 36 |
| M1O5J | 83adp_-1.03 | 31 | MBMLF | 59_-1.18 | 95 | MR5BT | 89adp_-1.29t | 87 |
| M1OEB | 96_-1.78t | 96 | MBZMQ | 86adp_-1.17t | 32 | MREGP | 12atp_-1.00 | 15 |
| M21ZT | 44atp_-0.87 | 13 | MC272 | 75adp_-1.22 | 99 | MRQ0O | 60_-1.19t | 34 |
| M2RXT | 57atp_-1.29t | 21 | MC3D0 | 41_-1.00t | 66 | MS1ZT | 5_-0.74 | 27 |
| M2TAA | 91_-1.67t | 91 | MC6BG | 5atp_-0.92 | 81 | MS4BB | 10atp_-0.93 | 40 |
| M2VPY | 92atp_-1.33t | 69 | MCPRP | 105atp_-1.11t | 54 | MSB6I | 92_-1.67t | 94 |
| M2VZK | 84atp_-1.26 | 20 | MCWDA | 8_-0.81 | 18 | MSCMP | 42_-1.01 | 46 |
| M30TD | 56atp_-1.01t | 90 | MDJWB | 52_-1.16 | 42 | MSPS6 | 58atp_-1.19t | 89 |
| M358F | 43atp_-0.65t | 41 | MDRU7 | 66_-1.28 | 77 | MSR51 | 102adp_-0.83 | 83 |
| M3DNA | 106atp_-1.20 | 55 | ME7LO | 78atp_-0.81t | 52 | MT56P | 2atp_-0.89t | 11 |
| M3NUL | 48_-1.11t | 58 | MEM9A | 1atp_-0.90t | 3 | MTQKH | 103_-2.41t | 105 |
| M3QQV | 87atp_-1.29 | 39 | MESSJ | 51atp_-1.00 | 6 | MTUAM | 85atp_-1.17t | 78 |
| M3RBZ | 94_-1.70t | 104 | MG4ML | 0atp_-1.16t | 88 | MTZB5 | 47atp_-0.87 | 28 |
| M3Y6D | 33_-1.00 | 33 | MGLVB | 53atp_-1.18 | 47 | MU4X7 | 77atp_-1.37t | 74 |
| M4GK5 | 54atp_-1.37 | 22 | MGRG1 | 97adp_-1.40t | 100 | MUCG2 | 104_-2.42t | 106 |
| M4RAZ | 45_-1.07t | 60 | MH9IN | 4atp_-0.95t | 51 | MUWA8 | 100adp_-1.25 | 43 |
| M4WKJ | 65_-1.27t | 97 | MHV6S | 99atp_-1.25 | 23 | MV1NF | 46atp_-1.13t | 37 |
| M50VT | 48atp_-0.64t | 61 | MIB32 | 38_-1.00t | 12 | MW0XQ | 9atp_-0.83t | 57 |
| M644A | 60atp_-0.89t | 14 | MIY6R | 20_-1.00t | 71 | MW80J | 49atp_-0.64 | 8 |
| M75CM | 67atp_-0.79t | 16 | MJ0WZ | 64_-1.27t | 50 | MWJWM | 3_-0.71t | 4 |
| M77O8 | 98adp_-1.40 | 68 | MJVZ1 | 82_-1.48 | 75 | MX2F5 | 96atp_-1.40t | 92 |
| M7QCU | 49_-1.12t | 53 | MKA7J | 14atp_-0.82 | 2 | MX31H | 74atp_-1.22 | 45 |
| M7U37 | 0_-0.58t | 5 | MKCPT | 11atp_-1.07t | 24 | MY02U | 89_-1.57t | 76 |
| M7ZLJ | 17_-1.00t | 19 | MKE6W | 73atp_-1.42t | 86 | MYCYM | 82atp_-1.03 | 65 |
| M839Z | 6atp_-0.73 | 30 | MKUGO | 93atp_-1.43t | 72 | MYLZR | 32_-1.00t | 73 |
| M8E3Q | 34atp_-0.86t | 26 | MKV9B | 50atp_-1.48 | 101 | MYOGH | 43_-1.04t | 80 |
| M8SCR | 80_-1.42 | 103 | ML01H | 91atp_-1.11t | 44 | MYV47 | 62atp_-1.36t | 1 |
| M8XF8 | 72_-1.35t | 48 | MMI5D | 56_-1.18t | 85 | MZ1F4 | 61_-1.21t | 38 |
| M8YL5 | 63atp_-1.10t | 56 | MN3VY | 88adp_-1.29t | 102 | MZH7E | 18_-1.00t | 10 |
| M929Q | 68atp_-1.03t | 29 | MNXBV | 93_-1.69t | 84 | MZSHY | 104adp_-1.49t | 98 |
| MAKA5 | 101atp_-0.83t | 67 | MNYAQ | 45atp_-0.55t | 17 |  |  |  |

**Table S7.** Thermostability.

| **Repo ID** | **ProtNLM** | **Length** | **Tm** | **Tm (2mM ATP/AMP-PNP)** |
| --- | --- | --- | --- | --- |
| M1X0B | Response regulator | 119 | 86.2 | n/a |
| MQLYM | Dihydroxyacetone kinase subunit L | 207 | 83.36 | n/a |
| MJFIU | Protein kinase domain-containing protein | 202 | 81.94 | n/a |
| MW51C | PAS domain S-box protein | 490 | 79.89 | n/a |
| M54FQ | LacI family transcriptional regulator | 296 | 76.8 | n/a |
| MV2G8 | Potassium transporter TrkA | 214 | 75.64 | n/a |
| M7PN6 | Ferritin | 159 | 71.05 | n/a |
| MTXPM | Trigger factor | 260 | 68.51 | n/a |
| MT47E | Toxin VapC | 127 | 67.05 | n/a |
| MLFIT | Ferritin | 157 | 65.18 | n/a |
| MWG7X | Prephenate dehydrogenase | 277 | 62.22 | n/a |
| MPCII | Hemerythrin domain-containing protein | 133 | 55.97 | n/a |
| MR1UH | 3-isopropylmalate dehydratase large subunit | 234 | 47.97 | n/a |
| M5CZB | 3.1.3.25 | 252 | 45.43 | n/a |
| MMUJG | Importin N-terminal domain-containing protein | 245 | 41.2 | n/a |
| MJY78 | DNA polymerase beta | 258 | 36.48 | n/a |
| M7S72 | Glycosyltransferase family 1 protein | 296 | 31.32 | n/a |
| M17H6 | AK | 189 | - | 58.4 |
| MS4BB | AK | 188 | - | 66.4 |
| MBMLF | AK | 214 | 53.1 | 55.5 |
| M21ZT | Sn-glycerol-3-phosphate ABC transporter ATP-bi | 357 | 62.9 | 63 |
| MEM9A | ABC transporter ATP-binding protein | 222 | 72.4 | 72.9 |
| MB5XC | ABC transporter ATP-binding protein | 234 | 83 | 83 |
| MREGP | Amino acid ABC transporter ATP-binding protein | 241 | 52.4 | 54.9 |
| MB11S | Amino acid ABC transporter ATP-binding protein | 233 | 67.6 | 71.4 |
| MSCMP | DEAD/DEAH box helicase | 296 | 77.2 | 76.9 |
| M3Y6D | DEAD/DEAH box helicase | 369 | 80.8 | - |
| MQZ1R | DEAD/DEAH box helicase | 296 | 52.5 | 53.8 |
| MKV9B | DEAD/DEAH box helicase | 299 | 56.4 | 57.1 |
| MUCG2 | DEAD/DEAH box helicase | 372 | 59.2 | 59.5 |
| M3RBZ | DEAD/DEAH box helicase | 372 | 66.5 | 66.1 |
| M1O5J | DEAD/DEAH box helicase | 295 | 73.8 | 73.8 |
| MPBDZ | ATP-dependent RNA helicase DbpA | 292 | 75.7 | 75 |
| M50VT | APS kinase | 197 | - | 38 |
| M3NUL | Adenylyl-sulfate kinase | 185 | 45.5 | 45.5 |
| MB6BK | AAA family ATPase | 280 | - | 66.4 |
| MSR51 | AAA domain-containing protein | 199 | 55.2 | 55.9 |
| M0VJA | Chromosome segregation protein SMC | 132 | 55.3 | 55.2 |
| M7QCU | Arsenical pump-driving ATPase | 240 | 65 | 67 |

## Notes

### Summary of Updates

Figures 1, 3, and 4 have been revised to include new data.

